# PASTA: Versatile Tyramine-oligonucleotide Amplification for Multi-modal Spatial Biology

**DOI:** 10.1101/2025.04.30.651463

**Authors:** Hendrik A Michel, Paige McCallum, Wenrui Wu, Jia Le Lee, Shuli Luo, Chi Ngai Chan, Johanna Schaffenrath, Yao Yu Yeo, Stephanie Pei Tung Yiu, Yang Wang, Lindsay Parmelee, Hongbo Wang, Melinda Burgess, Nourhan El Ahmar, Zoe Xiaozhu Zhang, Colm Keane, Tony Lim, Sabina Signoretti, Sonia Victoria Del Rincon, Bo Zhao, David R McIlwain, Yunhao Bai, Fei Chen, Sizun Jiang

## Abstract

Spatial proteomics techniques have revolutionized our understanding of tissue architecture, but are frequently limited by detection sensitivity, bioconjugation limitations, multiplexing capacity, and multi-modal integration. Here we present <u>P</u>rotein and nucleic <u>A</u>cid <u>S</u>erial <u>T</u>yramine <u>A</u>mplification (PASTA), a novel signal amplification approach that significantly enhances detection sensitivity while maintaining compatibility with diverse spatial profiling methodologies. PASTA utilizes horseradish peroxidase (HRP) recruitment pathways to generate tyramine radicals that deposit oligonucleotides, enabling adaptable signal amplification across multiple biomarkers at high-plex via cyclical imaging using complementary fluorophore-labeled oligonucleotides. We demonstrate that PASTA achieves up to 100-fold signal enhancement for markers with minimal background in blank controls. The method is compatible with *in situ* hybridization for DNA/RNA detection, proximity ligation assays for protein-protein interactions, sequential antibody staining protocols, or any modular combination thereof. PASTA enables antibody rescue of markers with suboptimal signal-to-noise ratios and is versatile in its applications to unconjugated antibodies, and multi-round probe-based RNA detection systems beyond current capabilities. This technique addresses key limitations in spatial-omics by enhancing sensitivity for challenging targets while maintaining compatibility with established multiplexing strategies, providing a versatile, cost-efficient, and valuable tool for comprehensive spatial tissue analysis in both research and clinical applications.

## Main

Spatial profiling technologies have revolutionized our understanding of tissue architecture by enabling visualization of biomolecules within their native microenvironment (1). Particularly, spatial proteomics has provided unprecedented insights into the complex cellular organization of tissues in health and disease, with high clinical utility (2). However, despite recent advances, several technical challenges continue to limit the comprehensive application of these technologies: 1) Low abundance targets often yield poor signal-to-noise ratios, 2) bioconjugation complications can restrict antibody and reagent availability, and 3) integrating multiple modalities remains complex and technically demanding.

Current spatial proteomics methods now routinely image > 20 antibody targets on the same tissue samples using cyclical imaging (3, 4) or lanthanide mass tags (5, 6), overcoming prior limitations in limited plex. Other challenges still remain, including sensitivity limitations, especially when detecting low-abundance markers critical for understanding immune responses and disease mechanisms. Signal amplification approaches like tyramide signal amplification (TSA) have been employed to enhance detection sensitivity but are limited in their multiplexing capabilities due to the biomolecules that can be deposited. Newer oligo-based amplification approaches, including SABER, are highly promising due to their high modular nature (7–10), but are currently limited to a single modality (i.e. only RNA or protein targets).

Here we present Protein and nucleic Acid Serial Tyramine Amplification (PASTA), a versatile signal amplification approach that significantly enhances detection sensitivity while maintaining compatibility with diverse spatial profiling methodologies. PASTA utilizes a wide range of horseradish peroxidase (HRP) recruitment pathways to generate tyramine radicals that covalently deposit oligonucleotides onto tissue sections (**Fig. 1A**). The PASTA method utilizes HRP-catalyzed oxidation of tyramine-oligonucleotide conjugates to generate phenoxy radicals that form stable carbon-carbon covalent bonds with electron-rich aromatic amino acids (primarily tyrosine residues) present in proteins around the biomolecule target of interest, resulting in permanently anchored oligonucleotide tags that resist degradation under stringent washing conditions and maintain functionality during long-term storage. Unlike traditional fluorophore-based approaches, the deposited oligonucleotides remain intact and functional for extended periods under proper storage conditions, allowing for delayed or repeated imaging sessions. This modular design not only enhances signal stability but also enables seamless integration with existing spatial technologies, while adding substantial signal amplification and multi-omics capabilities.

**Figure 1:**
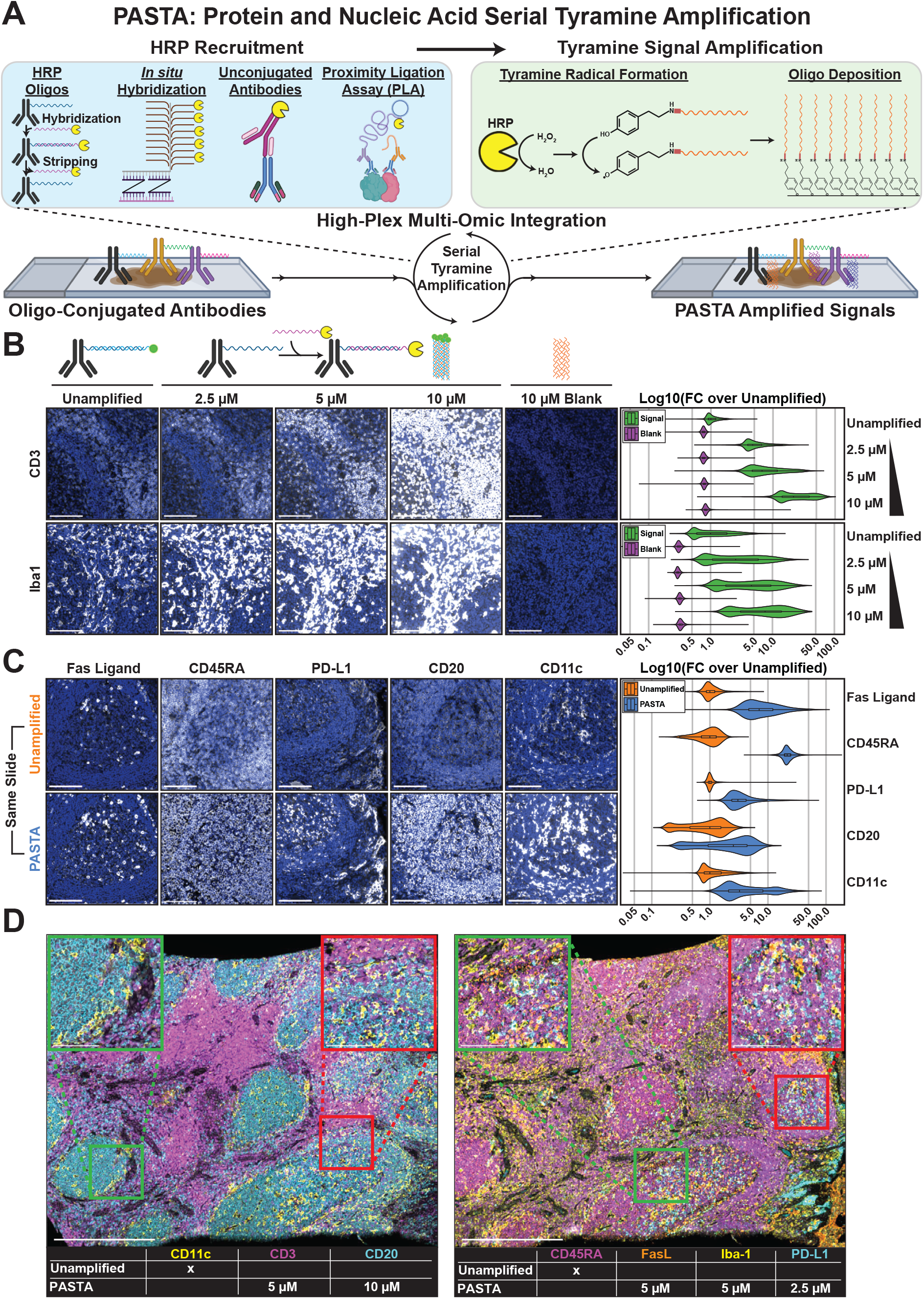
PASTA enables high-sensitivity spatial imaging with tunable amplification. **(A)** Overview schematic of the PASTA workflow. HRP recruitment including *in situ* hybridization, unconjugated antibodies, and proximity ligation assays enables tyramine radical formation. These radicals deposit oligos onto nearby proteins in the cells and tissues through covalent binding to tyrosine residues, creating stable protein-oligonucleotide conjugates that resist degradation and enable serial amplification. **(B)** Concentration-dependent signal amplification of CD3 and Iba1 in FFPE tissues. Images show comparison between unamplified detection (fluorescent oligonucleotide complementary to conjugated oligonucleotide) versus PASTA amplification at increasing tyramine-oligonucleotide concentrations. In the PASTA workflow, antibody-bound oligonucleotides are recognized by complementary oligo-HRP conjugates, which catalyze the deposition of distinct tyramine-oligonucleotide barcodes. Right panels show quantification of log10(fold change) in signal intensity compared to background controls, demonstrating concentration-dependent signal enhancement. Scale bars: 100 µm. **(C)** Comparison of unamplified versus PASTA-amplified detection for clinically relevant markers (Fas Ligand, CD45RA, PD-L1, CD20, CD11c) on the same tissue section. Violin plots quantify the significant signal enhancement achieved with PASTA amplification for each marker. Scale bars: 100 µm. **(D)** High-magnification imaging demonstrating spatial resolution and multiplexing capabilities. Left panel: CD11c (yellow), CD3 (magenta), and CD20 (cyan) with mixed amplification (unamplified CD11c, 5 µM PASTA for CD3, 10 µM PASTA for CD20). Right panel: CD45RA (magenta), FasL (yellow), Iba-1 (cyan), and PD-L1 (not colored) with varied PASTA amplification concentrations. Scale bars: 500 µm (main images), 100 µm (insets).

We integrated PASTA into the existing mature microfluidics platform, microscope, and image processing infrastructure of CODEX (3, 11), allowing repeated imaging through iterative stripping and hybridization of fluorophore-tagged complementary oligonucleotides. For PASTA amplification, we employed a sequential detection workflow: antibodies conjugated with specific oligonucleotide barcodes (i.e. barcode set A) were first bound and fixed to their targets, followed by recognition with oligo-HRP conjugates bearing complementary sequences to barcode set A. These HRP conjugates then catalyze the deposition of distinct tyramine-oligonucleotide barcodes (i.e. barcode set B) that differ from the initial antibody-conjugated sequences. These deposited barcodes are subsequently visualized using fluorophore-labeled oligonucleotides complementary to barcode set B. This strategic use of different barcode sets enables amplification while maintaining specificity. In contrast, the unamplified method directly visualized the antibody-bound oligonucleotides (barcode set A) with their respective fluorophore-labeled complements on the same tissue section. We demonstrate that this PASTA approach achieves up to 100-fold signal enhancement for critical immune markers in formalin-fixed paraffin-embedded (FFPE) clinically relevant tissues, including CD3 and Iba1, with minimal background in blank controls (imaged subsequent to chemical stripping of fluorescent oligonucleotide reporter) (**Fig. 1B, Fig. S1A**). The amplification effect scales with tyramine-oligonucleotide concentration, providing tunable sensitivity based on target abundance. A dual deposition experiment with a second PASTA deposited oligo with or without chemical stripping demonstrated that chemical stripping was sufficient for removal of HRP oligo activity (**Fig. S1B**). Further evaluation with clinically relevant markers including Fas Ligand, CD45RA, PD-L1, CD20, and CD11c confirmed substantial signal enhancement when comparing PASTA-amplified detection to unamplified direct detection on the same tissue section (**Fig. 1C**), demonstrating the compatibility of both detection methods within the same workflow. Importantly, PASTA maintains spatial resolution and signal specificity while significantly enhancing detection sensitivity, as evidenced by imaging of tonsil tissue sections with multiplexed markers of both amplified and unamplified channels (**Fig. 1D**).

Beyond enhancing standard antibody-based detection, PASTA demonstrates high versatility across multiple spatial biology applications. The method’s modular design enables integration with various spatial profiling techniques while maintaining high sensitivity and specificity.

PASTA seamlessly integrates with *in situ* hybridization approaches using PANINI (12) for viral nucleic acid detection, addressing the challenge of visualizing pathogens within their tissue context (**Fig. 2A**). We successfully applied PASTA to detect EBV episomal DNA in EBV-positive primary central nervous system lymphoma (PCNSL), and Spike RNA in SARS-CoV-2 infected rhesus macaque lung tissues. Staining for viral proteins and known negative tissues demonstrated that PASTA led to no loss of sensitivity or specificity (**Fig. S2A**). The tyramine-oligonucleotide deposition system amplified viral nucleic acid signals while enabling simultaneous phenotyping of surrounding cells through multiplex protein marker detection, including CD20, CD3, and CLDN5 for PCNSL, and CD20, S100A9, and Cytokeratin for SARS-CoV-2. This dual-modality approach allows correlative analysis between viral location and host cellular responses within the same tissue section, providing critical insights into infection biology (12).

**Figure 2:**
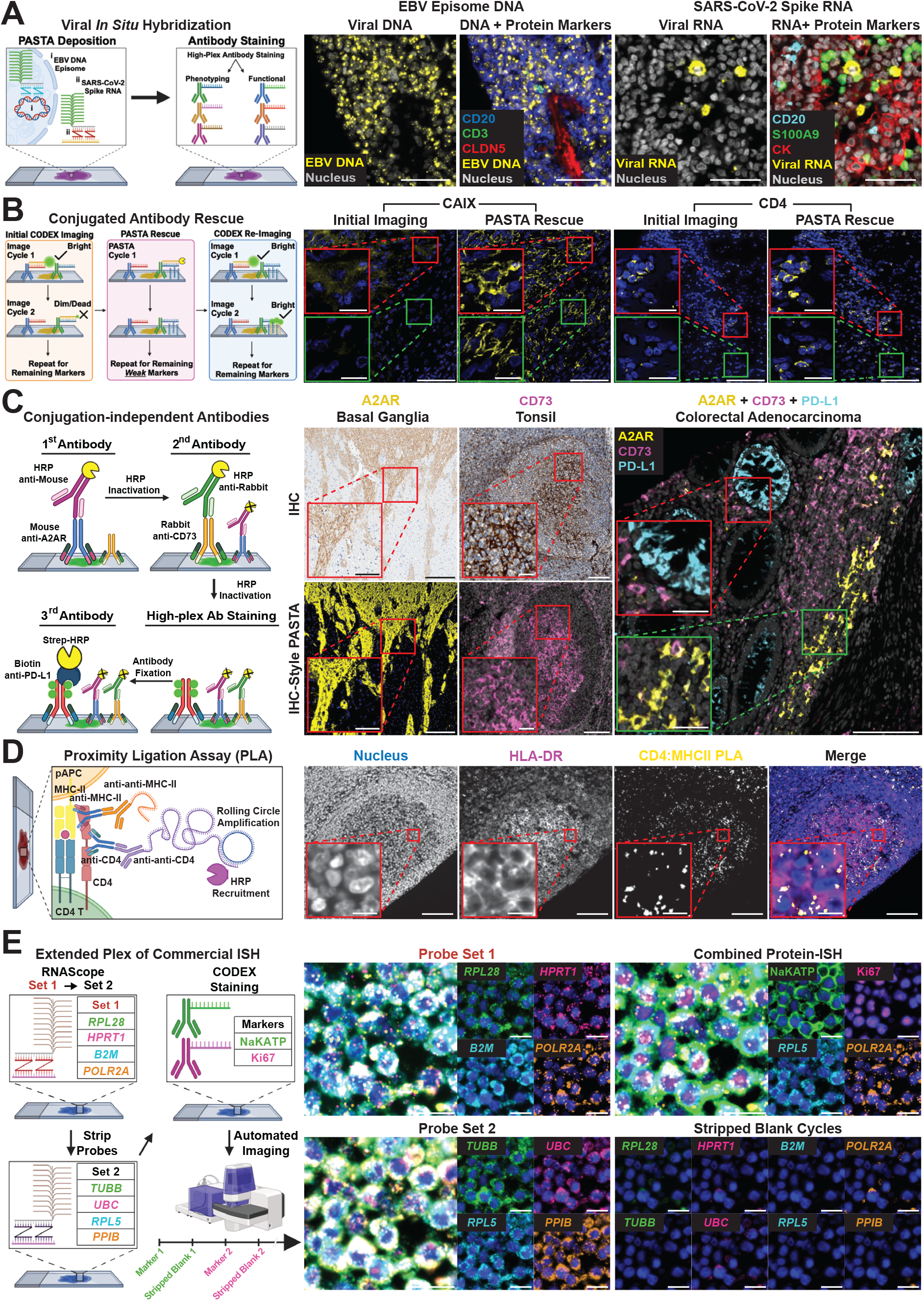
PASTA is highly versatile across multiple spatial biology applications. **(A)** Viral nucleic acid detection with PASTA. Left panels show EBV episomal DNA detection in PCNSL tissue (yellow), followed by simultaneous protein phenotyping (CD20 in blue, CD3 in magenta, CLDN5 in red). Right panels show SARS-CoV-2 spike RNA detection in infected rhesus macaque lung tissue (yellow) with protein marker co-staining (CD20, S100A9, Cytokeratin). PASTA enables strong nucleic acid signal amplification and protein visualization on the same tissue section through sequential application of *in situ* hybridization and antibody staining protocols. Scale bars: 50 µm. **(B)** Post hoc rescue of suboptimal markers using PASTA. Schematic (left) illustrates the workflow for conjugated antibody rescue through sequential PASTA cycles after initial imaging. For markers with insufficient signal intensity, tissues previously stained and imaged can undergo PASTA amplification, followed by reimaging. Center and right panels show real examples where initial imaging revealed weak signal for CAIX in tumor tissue and CD4 in a lymph node, which was dramatically enhanced after PASTA rescue amplification, enabling clear visualization of previously undetectable tissue structures and cellular localization. Scale bars: 100 µm (main images), 25 µm (insets). **(C)** Integration of unconjugated antibodies into multiplexed imaging. Sequential application workflow (left) shows how primary antibodies from different host species (mouse anti-A2AR, rabbit anti-CD73, biotinylated anti-PD-L1) are applied with HRP-conjugated secondary antibodies, with HRP inactivation between steps to prevent cross-reactivity. Comparative imaging of basal ganglia, tonsil, and colorectal adenocarcinoma demonstrates consistent staining patterns between traditional IHC (top row) and PASTA-enhanced IHC-style detection (bottom row), validating specificity while enabling higher multiplexing capability. Scale bars: 250 µm (A2AR main images), 100 µm (A2AR insets), 100 µm (other main images), 25 µm (other insets). **(D)** Proximity ligation assay (PLA) compatibility with PASTA and spatial proteomics. Schematic (left) shows the workflow for detecting CD4 interactions in antigen-presenting cells, including target recognition by primary antibodies, PLA probe binding, rolling circle amplification, and HRP recruitment for signal enhancement. Images show nuclear staining (blue), HLA-DR expression (magenta), CD4 PLA signal (yellow), and a merged view demonstrating the spatial relationships between these markers and the highly specific nature of the interaction detection. Scale bars: 100 µm (main images), 10 µm (insets). **(E)** Extended multiplexing of *in situ* hybridization using PASTA. Schematic (left) shows how two distinct RNA probe sets can be sequentially applied and amplified with PASTA on the same tissue section through probe stripping and reapplication in combination with multiplex immunofluorescence imaging for protein targets. Set 1: (*PPIB, TUBB, RPL5, UBC*) and Set 2: (*POLR2A, RPL28, B2M, HPRT1*) were applied followed by protein staining for the membrane marker NaKATP and the nuclear proliferation marker Ki67. Images show successful visualization of all 8 RNA targets with distinct spatial patterns on the same sample and in combination with protein targets, increasing the multiplexing capacity of standard off-the-shelf RNA ISH reagents. At the same time, the PASTA signal remained strippable during the automated imaging cycle as demonstrated using the blank cycles imaged at equal exposures. Scale bar: 25 µm.

A significant challenge in spatial proteomics is rescuing markers with suboptimal signal-to-noise ratios after the initial imaging data acquisition has been completed, especially for precious clinical samples. PASTA addresses this limitation through a post hoc amplification “rescue” that can be applied to tissues that have already undergone an initial round of spatial proteomics imaging (**Fig. 2B**). We demonstrate that markers such as CAIX (a critical tumor marker for clear cell renal cell carcinoma) and CD4 (a key immune marker that delineates CD4 T cells) that were barely detectable in initial imaging rounds exhibited significant signal boost when the same tissue sections were subsequently processed using PASTA amplification and reimaged. This capability allows users to return to previously imaged specimens, identify markers with insufficient signal post hoc, apply targeted amplification to those specific epitopes that are often critical to the study, and perform additional imaging sessions on these stained samples. This rescue approach salvages valuable spatial data without requiring new staining panels or additional clinical tissue sections, maximizing information yield from precious samples while allowing iterative improvement of spatial profiling results for meaningful biological insights. PASTA further addresses bioconjugation limitations by enabling incorporation of primary antibodies that may otherwise fail to be effectively labeled with biomolecules for detection, into highly multiplexed imaging workflows (**Fig. 2C**). We demonstrate this capability across diverse tissue types, including specificity assessment comparing PASTA amplified signals against traditional immunohistochemistry (IHC) in basal ganglia (A2AR), tonsil (CD73), and their combination in colorectal adenocarcinoma (A2AR, CD73, PD-L1). This approach uses sequential application of primary antibodies from different host species, followed by HRP-conjugated secondary antibodies that catalyze the deposition of distinct tyramine-oligonucleotide barcodes. Between antibody applications, HRP inactivation with H2O2 prevents cross-reactivity, allowing multiple unconjugated antibodies to be incorporated into the same workflow. Comparison between traditional immunohistochemistry (IHC) and PASTA-based detection (IHC-style PASTA) shows similar signal distributions and enables multiplexing capability by incorporation of PASTA with the CODEX spatial proteomics approach.

Protein-protein interactions are also critical for understanding cellular function but challenging to visualize *in situ*, with emerging translational insights (13). We demonstrate PASTA compatibility with proximity ligation assays (PLA) (**Fig. 2D**), enabling detection of protein interactions through PLA while maintaining compatibility with spatial proteomics. The method successfully detected CD4 interactions in antigen-presenting cells, with the amplified PLA signal (yellow) visible against nuclear (blue) and its binding partner HLA-DR (magenta) staining.

PASTA also enables extended multiplexing of commercial *in situ* hybridization (ISH) approaches (**Fig. 2E, Fig. S2B**). By applying serial probe stripping and re-probing protocols (see Material and Methods), we achieved visualization of two distinct probe sets (Set 1: *PPIB, TUBB, RPL5, UBC*; Set 2: *POLR2A, RPL28, B2M, HPRT1*) for a total of 8 RNA targets with PASTA signal amplification on the same FFPE cell pellet sections. The ability to strip probes completely and apply new ones without signal degradation dramatically increases the multiplexing capacity of standard ISH techniques on FFPE samples, whilst coupled with co-detection of protein targets, thus enabling robust spatial multi-omics in single tissue sections with standard microscopy setups and off-the-shelf reagents.

## Discussion

PASTA addresses critical challenges in spatial tissue profiling by providing a versatile signal amplification approach that maintains compatibility with diverse spatial biology modalities. The method’s ability to enhance detection of low-abundance targets while preserving multiplexing capabilities addresses a key limitation in current spatial proteomics approaches. By using directable HRP-catalyzed deposition of oligonucleotide barcodes, PASTA enables tunable signal amplification that can be adapted based on target abundance and specific application requirements.

The exceptional versatility of PASTA across multiple spatial biology domains represents a significant advancement for the field. We have demonstrated successful PASTA implementation on commercially available platforms and reagents including Akoya Biosciences’ PhenoCycler platform, ACDBio’s RNAscope reagents, and Na Vinci’s PLA kits. Importantly, the core PASTA chemistry is platform-agnostic and can be extended to virtually any spatial profiling technology or their combinations, making it broadly accessible to the research community using off-the-shelf reagents without requiring specialized equipment.

The compatibility of PASTA with existing commercial platforms makes it immediately accessible to researchers utilizing established spatial profiling workflows. The method’s integration with *in situ* hybridization, proximity ligation assays, and conventional antibody staining on the same tissue section enables true spatial multi-omics, allowing correlation between protein expression, nucleic acid detection, and biomolecular interactions. This multi-modal capability is particularly valuable for understanding complex tissue architectures in disease contexts where multiple analyte classes must be simultaneously visualized.

While some approaches for the quality control of multiplex imaging data have become available in recent years (14), suboptimal staining due to experimental limitations can be frustrating given the scarcity of the clinical samples or high associated costs. The post hoc amplification capability of PASTA for rescuing suboptimal markers represents a significant advancement in maximizing information yield from these precious clinical samples. This feature allows iterative improvement of spatial profiling results without requiring additional tissue sections or entirely new experimental setups, addressing a major challenge in translational research where sample availability is often limited.

PASTA’s application to unconjugated antibodies further expands the reagent options available for spatial profiling, allowing the incorporation of antibodies that may be resistant to conjugation chemistries while maintaining high-plex adaptability. The method’s demonstrated performance across diverse tissue types, including clinically relevant FFPE samples, combined with its adaptability to existing workflows and cost efficiency, makes PASTA a valuable tool for both research and clinical applications, particularly for investigations requiring the detection of low-abundance markers or complex spatial relationships in clinically relevant samples.

Future developments of the PASTA approach could include standardized reagent kits, optimization of additional detection modalities, and integration with automated staining platforms to enable reproducible staining with minimal human intervention. As spatial biology continues to advance toward clinical translation, PASTA provides a valuable tool to enhance detection sensitivity, provide resolutions for suboptimal results, and expand multi-modal capabilities, while maintaining compatibility with established workflows and instrumentation.

## Materials & Methods

### FFPE Tissues

The FFPE tissues used in this study were commercially available tonsil tissues (AMSBio, #AMS6022) for **Fig. 1** and **Fig. 2C** and commercially available HeLa cell pellets (ACDBio, #310045) for **Fig. 2E**. The basal ganglia (**Fig. 2C**) tissue was sectioned from a block of healthy, human, post-mortem basal ganglia which was a generous gift by Derek Allison (University of Kentucky). The remaining FFPE tissues were clinical samples used as part of ongoing studies. The PCNSL tissue (**Fig. 2A**) was part of a TMA generated by M.B. and C.K. The SARS-CoV-2-infected non-human primate tissues (15) (**Fig. 2A**) were a generous gift from Kristina De Paris (UNC, Chapel Hill) and Koen Van Rompay (UC Davis CNPRC). Renal cell carcinoma TMAs with tumor tissue (**Fig. 2B**, CAIX) and lymph nodes (**Fig. 2B**, CD4) were provided by N.E.A. and S.S. The colorectal adenocarcinoma tissue (**Fig. 2C**) was part of a TMA provided by Z.X.Z. and T.L.

### CODEX staining of FFPE tissues

FFPE slides were baked at 70°C for 1 hour and subsequently deparaffinized twice for 5 minutes each in Xylenes. To rehydrate the tissues, slides were mounted into a linear stainer (ST4020, Leica Biosystems) and incubated in the following solutions for 3 minutes each with dipping every 3 seconds: 3 x Xylenes, 2 × 100% Ethanol, 2 × 95% Ethanol, 1 × 80% Ethanol, 1 × 70% Ethanol, 3 x UltraPure Water (10977015, ThermoFisher). Slides were subsequently transferred to Dako pH 9 target retrieval solution (S2367, Agilent) preheated to 75°C in a PT module (ThermoFisher, A80400012), before being retrieved at 97°C for 20 minutes. After the PT module cooled down to 65°C, the slides were removed and cooled down on the bench for 10 minutes. Slides were then washed in UltraPure Water before the tissues were surrounded by a hydrophobic barrier pen (H-4000, Vector Laboratories). The pen was left to dry for 5 minutes while the tissue remained covered with UltraPure Water. The tissue was washed in UltraPure water before being covered with 3% H_2_O_2_ in 1X TBS (final, pH 7.5) and incubated for 10 minutes to inactivate endogenous peroxidases. After, the tissue was washed again in UltraPure Water before being washed in 1X TBS-T (0.05% Tween-20, pH 7.5). To block endogenous biotin, the tissue was covered with a few drops of avidin solution (927301, Biolegend) and incubated for 30 minutes before being washed thrice in 1X TBS-T. Then the tissue was covered with a few drops of biotin solution (927301, Biolegend) and incubated for 30 minutes before being washed thrice in 1X TBS-T.

To prepare the tissue for CODEX staining, the tissue was blocked with CODEX blocking solution (0.75X TBS-T, 3.75% Donkey Serum (D9663-10ML, Sigma), 0.75% Triton X-100, 0.0375% NaN_3_, 1 mg/ml sheared salmon sperm DNA (AM9680, ThermoFisher), 100 µg/ml mouse IgG (I5381-10mg, Sigma), 100 µg/ml rat IgG (I4141-10mg, Sigma), 100 nM of all unlabelled bottom oligos (Integrated DNA Technologies)) for 1 hour in a humidity chamber while photobleaching using strong LED lights (Best Buy, 6460231 and Amazon, B07C68N7PC). The temperature was monitored to not exceed 40°C. Meanwhile, the CODEX antibody cocktail was prepared as described previously(11). Briefly, conjugated antibodies (see **Table S1**) were spun down at 12,000 x g for 15 minutes at 4°C after which appropriate antibody volumes were added to antibody diluent (1X TBS-T, 5% Donkey Serum, 0.05% NaN_3_). The antibody mixture was then added to a 1X TBS-T pre-wetted 50 kDa centrifugal filter (UFC5050BK, Sigma) and spun down at 12,000 x g for 8 minutes. The concentrated antibody solution was eluted by inverting the filter into a fresh collection tube and spinning at 12,000 x g for 2 minutes. The volume of the solution was measured using a pipette. The volume needed to bring the volume of the solution to 0.75X of the final volume was added as antibody diluent to the filter and used to gently rinse through the filter. The filter was then eluted again by inverting it into the same collection tube and spinning again at 12,000 x g for 2 minutes. One quarter of the final volume was added as FFPE block (0.325X DPBS (ThermoFisher, 14190144), 162.5 mM NaCl, 25.35 mM NaH_2_PO_4_, 39.65 mM Na_2_HPO_4_, 1.625 mM EDTA, 0.1625% BSA, 0.02% NaN_3_, 0.05 mg/ml mouse IgG, 0.05 mg/ml rat IgG, 0.5 mg/ml sheared salmon sperm DNA, 100 nM of all unlabelled bottom oligos). Finally, the solution was filtered through a pre-wetted 0.1 µm filter (UFC30VV00, Sigma) by spinning at 12,000 x g for 2 minutes. After the blocking was completed, the excess blocking buffer was blotted off the slide and the antibody solution added to the tissues. The slides were left to incubate at 4°C overnight.

The next day, the slides were washed twice in S2 buffer (0.5X DPBS (14190144, ThermoFisher), 250 mM NaCl, 39 mM NaH_2_PO_4_, 61 mM Na_2_HPO_4_, 2.5 mM EDTA, 0.25% BSA, 0.02% NaN_3_) for 2 minutes each. Antibodies were fixed by incubating the tissues for 10 minutes in 1.6% paraformaldehyde (15710, Electron Microscopy Sciences) in S4 buffer (0.9X DPBS, 500 mM NaCl, 4.5 mM EDTA, 0.45% BSA, 0.02% NaN_3_) before being rinsed once and washed twice in 1X DPBS (14190144, ThermoFisher). Then the slides were submerged in ice-cold methanol on ice for 5 minutes before being rinsed once and washed twice in 1X DPBS. Finally, the tissues were treated with final fixative (4 µg/µl (from 200 µg/µl BS3 (21580, ThermoFisher) stock solution in DMSO) in 1X DPBS) for 20 minutes in the dark before being rinsed once and washed twice in 1X DPBS. The slides were stored in S4 buffer at 4°C until subsequent steps.

### PASTA amplification of CODEX antibody signal

To amplify the CODEX signal, slides were washed in azide-free 1X CODEX buffer (150 mM NaCl, 10 mM TrisHCl pH 7.5 (15567027, ThermoFisher), 10 mM MgCl_2_, 0.1% Triton X-100) twice for 5 minutes each. An initial chemical strip was performed to remove blocking oligos bound to the antibodies by incubating the slides twice in azide-free stripping buffer (80% DMSO, 20% azide-free 1X CODEX buffer) for 5 minutes each. Slides were then washed in azide-free 1X CODEX buffer. For each cycle of the serial tyramine deposition, the following steps were performed: First, the slides were incubated with HRP conjugated oligos or biotin conjugated oligos (see **Table S2**) at 200 nM in azide-free plate buffer (0.5 mg/ml sheared salmon sperm DNA (AM9680, ThermoFisher) in azide-free 1X CODEX buffer) for 20 minutes in the dark. In the case of biotin oligos, this was followed by two washes with azide-free 1X CODEX buffer and a 20-minute incubation with 1:100 HRP-Streptavidin (405210, Biolegend) in azide-free 1X CODEX buffer in the dark. Second, the slides were washed twice for 2 minutes each in azide-free 1X CODEX buffer and then once in 1X TBS for 2 minutes. Third, the tissues were incubated for 10 minutes with tyramine conjugated oligos diluted in TSA diluent buffer (FP1498, Akoya Biosciences) in the dark. Fourth, the slides were washed twice with azide-free 1X CODEX buffer. Finally, the HRP oligos were chemically stripped by incubating the slides twice for 5 minutes each with azide-free stripping buffer followed by two washes in azide-free 1X CODEX buffer for 2 minutes each. This process was repeated serially until all targets had been amplified.

### Testing HRP-oligo strippability for PASTA of CODEX antibody signal

In addition to the initial chemical strip with 2 × 5 minute washes in azide-free stripping buffer, the slides were incubated with 3% H_2_O_2_ in 1X TBS at 40°C for 30 minutes in a hybridization oven (ACDBio). Subsequent steps were carried out as outlined above until the end of the tyramine oligo incubation (here oligo 1), at which point the slides were washed twice for 2 minutes each in azide-free 1X CODEX buffer. The unstripped slide was moved to azide-free 1X CODEX buffer and stored in the dark until a later step. The stripped slide was incubated twice for 5 minutes each in azide-free stripping buffer followed by 2 washes for 2 minutes each in azide-free 1X CODEX buffer. At this point, the unstripped slide was retrieved and both slides were washed in 1X TBS for 2 minutes, after which the tissues were incubated with tyramine oligo (here oligo 2) diluted in TSA diluent buffer (FP1498, Akoya Biosciences) in the dark for 20 minutes. Finally, the slides were washed twice in azide-free 1X CODEX buffer for 2 minutes each. This step was repeated until all targets for strippability testing had been amplified.

### Automated Multiplex Imaging

For the Akoya PhenoCycler Fusion, slides were moved into 1X PBS from their storage solution (if applicable) to remove detergents. The edges of the slides were dried and a flow cell (Akoya Bioscience) was attached using the manufacturer’s flow cell press. The slides were subsequently incubated in 1X CODEX buffer for 10 minutes to allow the flow cell adhesives to harden. The PhenoCycler Fusion buffers were freshly prepared in-house and loaded into the machine. The slides were loaded into the machine following manufacturer instructions with an output setting for 16-bit images.

For the Akoya CODEX fluidics in combination with the Keyence BZ-X810 microscope, the machine was loaded with fresh 1X CODEX buffer and Dimethylsulfoxide. The coverslips were loaded into the fluidics stage and briefly stained with Hoechst 33342 (ThermoFisher, H3570) diluted 1:1,500 in plate buffer to identify imaging regions of interest. The automated machine was run following manufacturer’s instructions with an output setting for 16-bit images which were processed using the Akoya Singer software (v1.0.7).

Additionally, a reporter plate was prepared by adding ATTO550 or AlexaFluor647 conjugated oligos (GenScript, Biomers) complementary to the oligo barcodes on the antibodies or deposited on the tissue to a final concentration of 100 nM into plate buffer supplemented with Hoechst 33342 at a 1:300 dilution to a final volume of 250 µl. The wells were finally sealed with adhesive foil (ThermoFisher, AB0626).

### Image Quantification of CODEX SignalImage Quantification of CODEX Signal

Image Alignment: Due to the limited number of imaging cycles that can be carried out on the PhenoCycler Fusion at a time, the markers were imaged across four separate but sequential imaging runs. As a first step, the images from these runs were cropped into a single tonsil per image and subsequently aligned using VALIS (16) and combined into a single ome-tiff.

#### Image Segmentation and Feature Extraction

The images were segmented using MESMER (17) using default parameters and designating nuclear (Hoechst 33342) and membrane (conjugated oligo at 250ms exposure for CD3, CD11c, CD45RA, CD20, FasL, Iba1) markers. Single-cell feature extraction for the fluorescent signal for each cell was performed by summing up the pixel values for a marker for each cell and dividing it by the area of that cell.

#### Quantification and Data Presentation

For the markers of interest, a fold-change of the signals was calculated by dividing the signal of each cell by the median signal for the unamplified signal for all segmented cells in that specific tonsil tissue. For the unamplified signal in **Fig. 1B**, the signal for all three tissues was combined while for **Fig. 1C**, the unamplified signal was specifically for the same tissue condition as that shown for the PASTA signal. For the assessment of stripping sufficiency for HRP-oligo removal in **Fig. S1B**, the tissue treated with 10 µM tyramine oligonucleotides was analyzed by calculating the fold-change relative to the median signal for the unamplified signal of all segmented cells in that tissue.

### PASTA amplification for *in situ* hybridization of viral nucleic acids

FFPE slides stored at –20°C were removed from the freezer and allowed to warm to room temperature for 20-30 minutes. Subsequently, the slides were baked at 70°C for 1 hour and deparaffinized twice for 5 minutes each in Xylenes. To rehydrate the tissues, slides mounted into a linear stainer (ST4020, Leica Biosystems) and incubated in the following solutions for 3 minutes each with dipping every 3 seconds: 3 x Xylenes, 2 × 100% Ethanol, 2 × 95% Ethanol, 1 × 80% Ethanol, 1 × 70% Ethanol, 3 x Ultra-Pure Water (10977015, ThermoFisher). Slides were subsequently transferred to Dako pH 9 target retrieval solution (S2367, Agilent) preheated to 75°C in a PT module (ThermoFisher, A80400012), before being retrieved at 97°C for 20 minutes. After the PT module cooled down to 65°C, the slides were removed and cooled down on the bench for 10 minutes. Slides were then washed in UltraPure Water before the tissues were surrounded by a hydrophobic barrier pen (H-4000, Vector Laboratories). The pen was left to dry for 5 minutes while the tissue remained covered with UltraPure Water. The tissue was washed in UltraPure water before being covered with RNAScope Hydrogen Peroxide (ACDBio, #322335) and incubated for 10 minutes at room temperature (SARS-CoV-2 RNA) or 20 minutes at 40°C (EBV DNA) to block endogenous peroxidase activity. After, the tissue was washed again in UltraPure Water. C2-C4 probes were diluted 1:50 in C1 probe solution (see **Table S3**) and then prewarmed for an additional 15 minutes. 50 µl of the probe solution was added to the tissues before being placed in a humidity tray which was placed in a hybridization oven (ACDBio) overnight at 40°C.

The next day, the slides were washed twice for 2 minutes each in 0.5X RNAScope Wash Buffer (ACDBio, #310091). Next, 2 drops of Multiplex v2 Amp 1 (ACDBio, #323101) were added to each tissue and incubated for 30 minutes at 40°C. The slides were rinsed and washed twice followed by incubation with 2 drops of Multiplex v2 Amp 2 (ACDBio, #323102) for 15 minutes at 40°C. The slides were rinsed and washed twice followed by incubation with 2 drops of Multiplex v2 Amp 3 (ACDBio, #323103) for 30 minutes at 40°C. The slides were rinsed and washed twice followed by incubation with 2 drops of Multiplex v2 HRP-C1 (ACDBio, #323104) for 15 minutes at 40°C. The slides were rinsed and washed twice followed by incubation with 2 drops of RNAScope Brown Kit Amp 5 (ACD-Bio, #322315) for 45 minutes at room temperature. The slides were rinsed and washed twice in 1X TBS-T followed by incubation with 2 drops of RNAScope Brown Kit Amp 6 (ACDBio, #322316) for 15 minutes at room temperature. The slides were rinsed and washed twice followed by incubation with tyramine oligo diluted to a final concentration of 5 µM in TSA diluent buffer (FP1498, Akoya Biosciences) at room temperature for 15 minutes. After washing again, 2 drops of HRP Blocker (ACDBio, #323107) were incubated for 15 minutes at 40°C followed by 2 washes. This process was repeated for the remaining channels as applicable. Following the full RNAScope protocol, the tissues were washed with 1X TBS-T twice for 2 minutes each. Then tissues were blocked and stained with CODEX antibodies as described above and incubated overnight at 4°C. The next day, the tissues were washed and fixed as described above. The slides were imaged on the PhenoCycler Fusion.

### Rescue of conjugated antibody signal using PASTA

Following the identification of markers requiring signal amplification for marker rescue, the coverslips were rinsed twice for 2 minute each in 1X TBS-T. As the tissues had previously been stained with a biotinylated antibody and treated with fluorescent streptavidin, an additional avidin-biotin block was carried out by incubating 20 minutes with avidin solution (927301, Biolegend), washing twice for 2 minutes each in 1X TBS-T, incubating 20 minutes with biotin solution (927301, Biolegend) and finally washed twice for 2 minutes each in 1X TBS-T. To block the activity of previous HRP conjugated antibodies, the tissues were incubated with HRP Blocker (ACDBio, #323107) for 15 minutes at 40°C followed by 2 washes for 2 minutes each in 1X TBS-T.

Subsequently, the tissues were stripped, washed, incubated with 100 nM biotin or HRP oligos (see **Table S2**) in azide-free plate buffer for 10 minutes in the dark, washed, stained with streptavidin HRP (ThermoFisher, S911) diluted 1:100 in azide-free 1X CODEX buffer for 20 minutes in the dark, washed, incubated with 3 µM of tyramine oligo diluted in TSA diluent buffer (FP1498, Akoya Biosciences) for 10 minutes, and finally washed again. The process was carried out as above and repeated for all markers to be rescued. The coverslips were reimaged in an automated run on the Keyence microscope with CODEX fluidics. The rescued markers were imaged using AlexaFluor488 oligos (see **Table S1**).

### Integration of unconjugated and biotinylated antibodies into CODEX using PASTA

Deparaffinization and antigen retrieval were performed on study coverslips using conditions described above. After the retrieved coverslips have been cooled to room temperature, they were washed in UltraPure Water before the tissues were surrounded by a hydrophobic barrier pen (H-4000, Vector Laboratories). The pen was left to dry for 5 minutes while the tissue remained covered with UltraPure Water. The tissues were then rinsed three times with 1X TBS-T (0.05% Tween-20, pH 7.5) for 2 minutes each. To block endogenous biotin, the tissues were covered with a few drops of avidin solution (927301, Biolegend) and incubated for 20 minutes, with avidin being refreshed at the 10 minute mark, before being washed thrice in 1X TBS-T. The tissues were then covered with a few drops of biotin solution (927301, Biolegend) and incubated for 20 minutes, with biotin being refreshed at the 10 minute mark, before being washed thrice in 1X TBS-T. Following avidin and biotin blocking, the tissues were blocked with blocking solution (0.75X TBS-T, 3.75% Donkey Serum (D9663-10ML, Sigma), 0.75% Triton X-100, 0.0375% NaN_3_) for 1 hour at room temperature. The primary unconjugated antibodies (see **Table S1**) were diluted in antibody diluent (1X TBS-T, 5% Donkey Serum, 0.05% NaN_3_) and added to the tissues following the blocking. The slides were incubated overnight at 4°C.

The next day, the tissues were washed three times with 1X TBS-T (0.05% Tween-20, pH 7.5) for 2 minutes each. The tissues were then covered with 3% H_2_O_2_ in 1X TBS (final, pH 7.5) and incubated for 15 minutes at room temperature to inactivate endogenous peroxidases. After, the tissues were washed thrice with 1X TBS-T (0.05% Tween-20, pH 7.5) before the tissues were incubated with the first secondary antibody; anti-mouse HRP (Origene, #D37-110) to target the primary antibody derived from a mouse host for 30 minutes at room temperature. The tissues were then washed thrice with 1X TBS-T (0.05% Tween-20, pH 7.5) before being incubated with tyramine conjugated oligos in TSA diluent buffer (FP1498, Akoya Biosciences) in the dark for 15 minutes. The tissues were again washed with 1X TBS-T (0.05% Tween-20, pH 7.5). To quench the active HRP activity, the coverslips were incubated with 3% H_2_O_2_ in 1X TBS (final, pH 7.5) for 30 minutes at 40°C. After rinsing the coverslips thrice with 1X TBS-T (0.05% Tween-20, pH 7.5), the tissues were incubated with the second secondary antibody; anti-rabbit HRP (Origene, #D13-110) to target the primary antibody derived from a rabbit host for 30 minutes at room temperature. The tissues were then washed thrice with 1X TBS-T (0.05% Tween-20, pH 7.5) before being incubated with tyramine conjugated oligos in TSA diluent buffer (FP1498, Akoya Biosciences) (note: the oligo must be distinct from that from the first round) in the dark for 15 minutes. The tissues were again washed with 1X TBS-T (0.05% Tween-20, pH 7.5). Active HRP activity was again quenched by incubating the coverslips with 3% H_2_O_2_ in 1X TBS (final, pH 7.5) for 30 minutes at 40°C. The tissues were then washed thrice with 1X TBS-T (0.05% Tween-20, pH 7.5) to remove any residual H_2_O_2_.

After the tyramine oligos were deposited, the tissues were incubated with CODEX blocking solution (0.75X TBS-T, 3.75% Donkey Serum (D9663-10ML, Sigma), 0.75% Triton X-100, 0.0375% NaN_3_, 1 mg/ml sheared salmon sperm DNA (AM9680, ThermoFisher), 100 µg/ml mouse IgG (I5381-10mg, Sigma), 100 µg/ml rat IgG (I4141-10mg, Sigma), 100 nM of all 58 unlabelled bottom oligos (Integrated DNA Technologies)) and allowed to incubate while photobleaching for 1 hour. CODEX antibodies including the biotinylated antibody were prepared as described above and incubated overnight at 4°C. After fixation as described above, the tissues were washed with 1X TBS-T (0.05% Tween-20, pH 7.5) and then incubated with HRP-Streptavidin (S911, ThermoFisher) diluted 1:100 in 1X TBS-T (0.05% Tween-20, pH 7.5) for 30 min at room temperature. Subsequently, the slides were washed with 1X TBS-T (0.05% Tween-20, pH 7.5) before being incubated with tyramine oligos in TSA Diluent in the dark for 15 minutes. Finally, the slides were washed twice with 1X TBS-T (0.05% Tween-20, pH 7.5) and twice with azide-free 1X CODEX buffer (150 mM NaCl, 10 mM TrisHCl pH 7.5 (15567027, ThermoFisher), 10 mM MgCl_2_, 0.1% Triton X-100) twice for 5 minutes each.

### PASTA amplification of proximity ligation assay (PLA) signal

Deparaffinization and antigen retrieval were performed on study slides using conditions described above. After the retrieved slides have been cooled to room temperature for 20 minutes, the slides were washed once in 1X TBS-T for 5 minutes followed by S2 buffer for 20 minutes after which the tissues were circled with a hydrophobic barrier pen. Then the tissues were blocked with blocking buffer (PLA Buffer (Navinci), supplemented with 0.5 mg/ml sheared salmon sperm DNA and 25 nM of unlabelled bottom oligos) for 1 hour while photobleaching.

The PLA reaction was carried out using a modified version of the manufacturer’s protocol. Briefly, primary antibodies were incubated at 1:100 dilution in Navenibody diluent (Navinci) at 37°C for 1 hour. The diluted navenibodies M1 and R2 (Navinci) were diluted to 1X working concentration in navenibody diluent (Navinci) and added to the tissues followed by 1 hour incubation at 37°C. The slides were rinsed once and washed twice for 5 minutes each in 1X TBS-T prewarmed to 37°C. Naveni Buffer 1 and Naveni Enzyme 1 (Navinci) were diluted in UltraPure Water to 1X working concentration and incubated on the tissues for 30 minutes at 37°C. The slides were rinsed once and washed once in 1X TBS-T. Naveni Buffer 2 and Naveni Enzyme 2 (Navinci) were diluted in UltraPure Water to 1X working concentration and subsequently incubated on the tissues for 90 minutes at 37°C. The slides were washed twice for 5 minutes each in 1X TBS before HRP Reagent (Navinci) diluted in HRP Diluent (Navinci) was incubated for 30 minutes at room temperature. Finally, the slides were washed twice for minutes each in 1X TBS. For the TSA reaction, the tyramine conjugated oligo and tyramine conjugated digoxigenin were diluted to 5 µM and 1:250, respectively, in TSA Diluent (Akoya) and incubated on the tissue for 10 minutes at room temperature. Slides were washed twice for 5 minutes each in 1X TBS. CODEX antibody cocktail was prepared as described above, added to the slides and incubated overnight at 4°C. Antibodies were fixed as described above and slides were imaged on the PhenoCycler Fusion.

### Combined High-plex RNAScope *in situ* hybridization with antibody staining using PASTA

FFPE slides were baked at 70°C for 1 hour and subsequently deparaffinized twice for 5 minutes each in Xylenes. To rehydrate the tissues, slides mounted into a linear stainer (ST4020, Leica Biosystems) and incubated in the following solutions for 3 minutes each with dipping every 3 seconds: 3 x Xylenes, 2 × 100% Ethanol, 2 × 95% Ethanol, 1 × 80% Ethanol, 1 × 70% Ethanol, 3 x UltraPure Water (10977015, ThermoFisher). Slides were subsequently transferred to Dako pH 9 target retrieval solution (S2367, Agilent) preheated to 75°C, before being retrieved at 97°C for 20 minutes. After the PT module cooled down to 65°C, the slides were removed and cooled down on the bench for 10 minutes. Slides were then washed in UltraPure Water before the tissues were surrounded by a hydrophobic barrier pen (H-4000, Vector Laboratories). The pen was left to dry for 5 minutes while the tissue remained covered with UltraPure Water. The tissue was washed in UltraPure water before being covered with RNAScope Hydrogen Peroxide (ACDBio) and incubated for 10 minutes at room temperature to block endogenous peroxidase activity. After, the tissue was washed again in UltraPure Water. C2-C4 probes were diluted 1:50 in C1 probe solution (see **Table S3**) and then prewarmed for an additional 15 minutes. 50 µl of the probe solution was added to the tissues before being placed in a humidity tray which was placed in a hybridization oven (ACDBio) overnight at 40°C.

The next day, the slides were rinsed once and washed twice for 2 minutes each in 0.5X RNAScope Wash Buffer. 3 drops of prewarmed Multiplex v2 Amp 1 were added and incubated for 30 minutes at 40°C. Next, the slides were rinsed and washed as above followed by addition of 3 drops of prewarmed Multiplex v2 Amp 2 and incubated for 30 minutes at 40°C. The slides were rinsed and washed again followed by addition of 3 drops of prewarmed Multiplex v2 Amp 3 and incubated for 15 minutes at 40°C. For the development of the 4 multiplex channels (C1-C4), the following steps were repeated for each channel: 3 drops of HRP-C_n_ were added to each slide and incubated for 15 minutes at 40°C. This was followed by a rinse and two washes. Meanwhile, tyramine oligo or fluorophore (for 4-channel positive control, see **Table S3**) were diluted in TSA diluent buffer (FP1498, Akoya Biosciences) to a concentration of 5 µM (tyramine oligo) or 1:100 (tyramine fluorophores) and added to the tissue for incubation at room temperature for 20 minutes in the dark. Slides were rinsed and washed twice, after which 3 drops of HRP blocker were added to each tissue and incubated at 40°C for 15 minutes followed by a final rinse and wash. For the fluorophore controls, the following tyramine conjugates were deposited in order (see **Table S3**): digoxigenin tyramine (Akoya Biosciences, #NEL748001KT), AlexaFluor647 tyramine (ThermoFisher, #B40958), AlexaFluor568 tyramine (ThermoFisher, #B40956), and AlexaFluor 488 tyramine (ThermoFisher, #B40953).

For the fluorophore control slides, the slides were washed in 1X TBS-T for 2 minutes after the final RNAScope deposition. The tissues were blocked with blocking solution (0.75X TBS-T, 3.75% Donkey Serum (D9663-10ML, Sigma), 0.75% Triton X-100, 0.0375% NaN_3_) for 1 hour at room temperature followed by staining with mouse anti-digoxigenin antibody (Novus Biologicals, #NBP2-31191) diluted 1:500 in antibody diluent (1X TBS-T, 5% Donkey Serum, 0.05% NaN_3_) for 1 hour at room temperature. After the incubation, the slides were washed twice with 1X TBS-T for 2 minutes each followed by secondary antibody staining with anti-mouse DyLight755 (ThermoFisher, #SA5-10175) diluted 1:100 in antibody diluent and incubated for 30 minutes at room temperature. The slides were washed twice with 1X TBS-T for 2 minutes each and finally stained with Hoechst 33342 (ThermoFisher, H3570) diluted in 1X TBS-T followed by two washes in 1X TBS-T for 2 minutes each. Slides were coverslipped with Prolong Gold (ThermoFisher, #P36930) and allowed to dry before being imaged with the PhenoImager Fusion (Akoya Bio-science).

After all 4 channels had been developed, the slides were placed in prewarmed stripping solution (90% Dimethylsulfoxide, 10% 0.5X RNAScope Wash Buffer) twice for 10 minutes each at 42°C. Slides were washed twice in 0.5X RNAScope Wash Buffer and then placed in UltraPure water. Meanwhile, C2-C4 probes were diluted 1:50 in C1 probe solution and then prewarmed for an additional 15 minutes. 50 µl of the probe solution was added to the tissues before being placed in a humidity tray which was placed in the hybridization oven overnight at 40°C. The next day, the channels were developed as described above until all RNAScope markers had been developed. Slides were not stripped after the last RNAScope iteration.

To combine antibody staining, slides were washed in 1X TBS-T for 5 minutes twice. Then the tissues were incubated with CODEX blocking solution and blocked for 2 hours while photobleaching as described above. The tissues were then incubated with an antibody cocktail overnight at 4°C. The next day, the tissues were processed and fixed as described above and stored in S4 buffer until imaging on the PhenoCycler Fusion. For increased sensitivity, the raw images (stitched but not aligned or background subtracted) were aligned using VALIS(16) and combined into a single ome-tiff.

## DATA AVAILABILITY

The data in this manuscript will be made available on Zenodo upon publication.

## CODE AVAILABILITY

All code used in this manuscript for the plotting of quantified images will be made available in a github repository upon publication.

## ACKNOWLEDGEMENTS

We acknowledge the Akoya Biosciences support team, especially Craig Lassy and Michael Hair, for their technical support, GenScript, Biomers, and Integrated DNA Technologies for their development of tyramine and HRP oligos, Na Vinci and ACD-Bio for their generous donation of reagents utilized in this study, and Parhelia Biosciences for equipment support for automated slide staining on the OpenTrons OT-2. We also thank all members of the Jiang Lab for fruitful discussions and support of this project.

We thank Kristina De Paris and Koen Van Rompay for providing SARS-CoV-2-infected NHP tissues. We thank Derek Allison (University of Kentucky) for providing basal ganglia FFPE tissues.

S.J. is supported in part by NIH DP2AI171139, P01AI177687, R01AI149672, a Gilead’s Research Scholars Program in Hematologic Malignancies, the Bill & Melinda Gates Foundation INV-002704, the Dye Family Foundation, and the Bridge Project, a partnership between the Koch Institute for Integrative Cancer Research at MIT and the Dana-Farber/Harvard Cancer Center. J.L.L. is supported by a National Science Scholarship (PhD) from the Agency for Science, Technology and Research, Singapore (BM/NDR/18/003) and is a Schmidt Science Fellow. S.P.T.Y is a MacMillan Family Foundation Awardee of the Life Sciences Research Foundation. Y.Y.Y. is a recipient of the Albert J Ryan Fellowship. J.S. is supported by a Roche Postdoctoral Fellowship.

S.V.D.R. is supported by the Canadian Cancer Society Grant 707140. P.M. is supported by the Canadian Institute of Health Research Grant 195177 and the Graduate Research Enhancement and Travel Award. S.S. is supported by the NCI Dana Farber / Harvard Cancer Center Kidney Cancer SPORE grant (P50-CA101942-12). Schematics in Figures 1 and 2 were generated using Biorender.

This article reflects the views of the authors and should not be construed as representing the views or policies of the institutions that provided funding.

## AUTHOR CONTRIBUTIONS

Conceptualization: S.J., H.A.M

Experiment: H.A.M., P.M., J.L.L., S.L., C.N.C., J.B.S., Y.Y.Y., S.P.T.Y., L.P., H.W. Analysis: H.A.M.

Contribution of unique reagents, tools, or technical expertise: W.W., B.Z., Y.B., F.C., M.B., C.K., N.E.A., S.S., Z.X.Z., T.L., S.V.D.R., D.R.M.

Writing: H.A.M., S.J. Supervision and funding: S.J.

