## Supplemental Figures and Tables for "PASTA: Versatile Tyramine-oligonucleotide Amplification for Multi-modal Spatial Biology"

[1]Center for Virology and Vaccine Research, Beth Israel Deaconess Medical Center, Harvard Medical School, Boston, MA, United States
[2]Program in Virology, Division of Medical Sciences, Harvard Medical School, Boston, MA, United States
[3]Division of Experimental Medicine, McGill University, Montreal, QC, Canada
[4]Division of Infectious Diseases, Department of Medicine, Brigham and Women's Hospital, Harvard Medical School, Boston, Massachusetts, USA
[5]Frazer Institute, Translational Research Institute, University of Queensland, Brisbane, Queensland, Australia
[6]Department of Pathology, Brigham and Women's Hospital, Boston, MA, United States
[7]Department of Anatomical Pathology, Singapore General Hospital, Singapore, Singapore
[8]Department of Microbiology and Immunology, University of Nevada Reno, Reno, NV, USA
[9]Broad Institute of Harvard and MIT, Cambridge, MA, United States

### Supplementary Figures

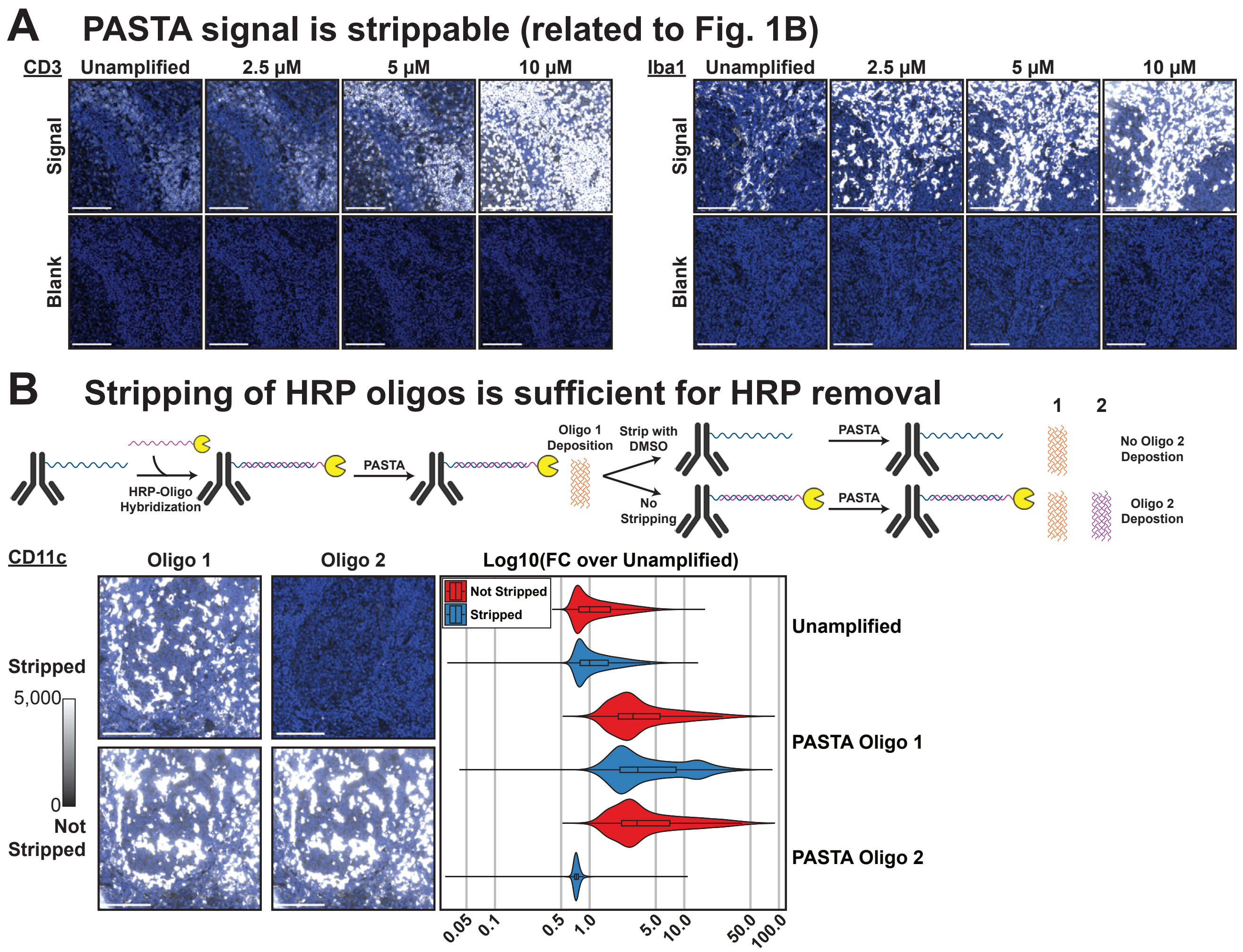

**Supplementary Figure 1: PASTA is a robust approach and allows reduced imaging time. (A)** PASTA amplification and related blank cycles related to data shown in **Fig. 1B**. The blank cycles are in the cycle immediately following the signal acquisition at equal exposure. Equal minimum and maximum pixel representation for the PASTA amplified image for each marker and corresponding blanks. Scale bars: 100μm. **(B)** Chemical stripping of HRP oligos is sufficient to remove HRP activity. Schematic (top) shows previously PASTA amplification as introduced in **Fig. 1** for deposition of oligo 1 followed by treatment with DMSO to chemically strip HRP-oligo or lack thereof before carrying out deposition of oligo 2. As expected, chemical stripping is sufficient to reduce oligo B signal to below unamplified signal thus demonstrating its sufficiency for HRP activity removal. Plot shows log10 of the fold-change relative to the unamplified signal. Scale bar: 100μm.

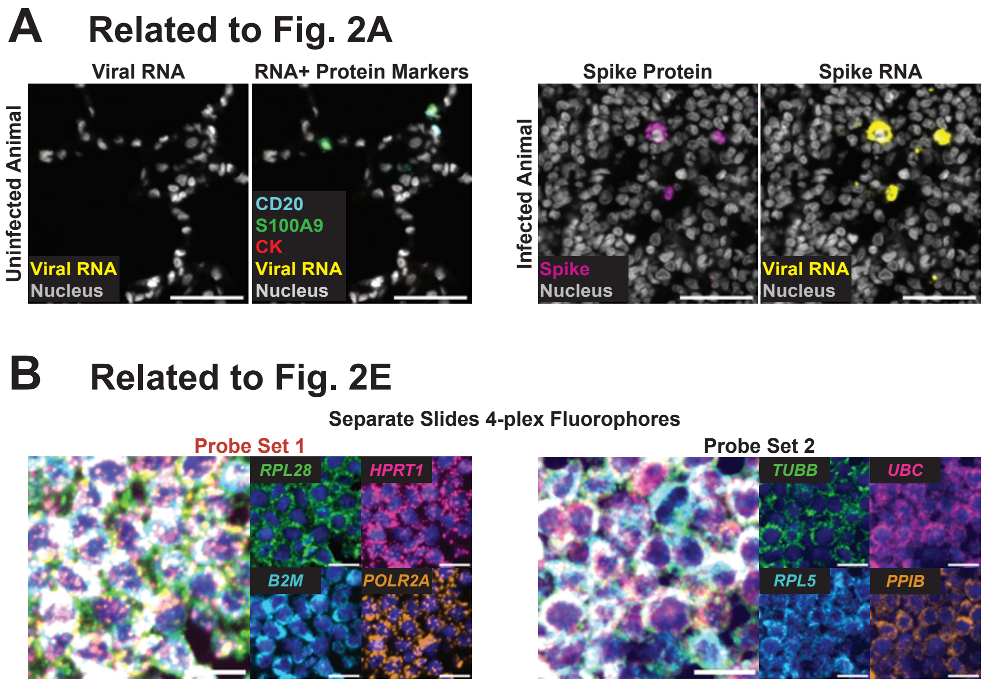

**Supplementary Figure 2: Integration of multiple spatial platforms with PASTA is robust.** **(A)** SARS-CoV-2 viral RNA detection related to **Fig. 2A**. No viral RNA and few infiltrating immune cells were detected in the uninfected animal tissue (left). In infected animals, cells positive for viral RNA were also positive for the spike protein (right). Scale bar: 50µm. **(B)** Robustness of the extended multiplexing using PASTA related to **Fig. 2E**. Left: The two probe sets were used separately with fluorophores to confirm the expected staining. Scale bar: 25µm.

### Supplementary Tables

| Figure 1 | | | | | | | |
|---|---|---|---|---|---|---|---|
| **Target** | **Clone** | **Vendor** | **Catalog number** | **Oligo** | **Fluorophore** | **Titer (dilution or ug/ml)** | **Exposure (ms)** |
| Iba1 | E4O4W | CST | 60083 | 20 | AlexaFluor-647 | 2.5 | 250ms, 50ms, 10ms |
| CD45RA | HI100 | Biolegend | 304102 | 72 | AlexaFluor-647 | 1:150 | 250ms, 50ms, 10ms |
| PD-L1 | E1L3N | CST | 85164SF | 11 | AlexaFluor-647 | 1 | 250ms, 50ms, 10ms |
| CD20 | E7B7T | CST | 34154BC | 48 | AlexaFluor-647 | 2 | 250ms, 50ms, 10ms |
| CD3 | D7A6E | CST | 71928BC | 77 | AlexaFluor-647 | 5 | 250ms, 50ms, 10ms |
| Tox/Tox2 | E6I3Q | CST | 62886SF | 74 | AlexaFluor-647 | 5 | 250ms, 50ms, 10ms |
| CD11c | EP1347Y | Abcam | ab216655 | 49 | AlexaFluor-647 | 2.5 | 250ms, 50ms, 10ms |
| Fas Ligand | polyclonal | Abcam | ab134401 | 7 | AlexaFluor-647 | 5 | 250ms, 50ms, 10ms |
| **Figure 2A: EBV DNA Detection in PCNSL** | | | | | | | |
| **Target** | **Clone** | **Vendor** | **Catalog number** | **Oligo** | **Fluorophore** | **Titer (dilution or ug/ml)** | **Exposure (ms)** |
| CD20 | E7B7T | CST | 34154 | 48 | ATTO550 | 5 | 150ms |
| CD3e | D7A6E | CST | 71928BC | 77 | ATTO550 | 2.5 | 150ms |
| CD3 | MRQ-39 | CellMarque | 103R-200UG | 77 | ATTO550 | 2.5 | 150ms |
| CLDN5 | E5D9Y | CST | 78328 | 49 | AlexaFluor-647 | 5 | 150ms |
| **Figure 2A: SARS-CoV-2 Spike RNA Detection in infected non-human primates** | | | | | | | |
| **Target** | **Clone** | **Vendor** | **Catalog number** | **Oligo** | **Fluorophore** | **Titer (dilution or ug/ml)** | **Exposure (ms)** |
| CD20 | rIGEL/773 | Novus | NBP2-54591-200ug | 48 | ATTO550 | 1:100 | 50ms |
| S100A9 | MAC387 | GeneTex | GTX76579 | 14 | ATTO550 | 1:50 | 50ms |
| Cytokeratin | C11 | Biolegend | 628602 | 67 | AlexaFluor-647 | 1:50 | 50ms |
| **Figure 2B: Rescue of markers in RCC samples** | | | | | | | |
| **Target** | **Clone** | **Vendor** | **Catalog number** | **Oligo** | **Fluorophore** | **Titer (dilution or ug/ml)** | **Exposure (ms)** |
| CAIX | EPR23055-5 | Abcam | ab270401 | 62 | AlexaFluor-647 (original)<br>AlexaFluor-488 (rescue) | 10 | 500ms (original)<br>800ms (rescue) |
| CD4 | EPR6855 | Abcam | ab181724 | 20 | ATTO550 (original)<br>AlexaFluor-488 (rescue) | 8 | 250ms (original)<br>1000ms (rescue) |
| **Figure 2C: Conjugation-independent integration in colorectal carcinoma samples** | | | | | | | |
| **Target** | **Clone** | **Vendor** | **Catalog number** | **Oligo** | **Fluorophore** | **Titer (dilution or ug/ml)** | **Exposure (ms)** |
| CD73 | D7F9A | CST | 87661 | uncon. | AlexaFluor-647 (tonsil)<br>AlexaFluor-750 (CRC) | 0.1 | 200ms (tonsil)<br>300ms (CRC) |
| A2AR | 7F6-G5-A2 | Abcam | ab79714 | uncon. | ATTO550 | 0.5 | 250ms |
| PD-L1 | E1L3N | CST | 85164SF | Biotin | AlexaFluor-647 | 5 | 50ms |
| **Figure 2D: Integration of proximity ligation assays** | | | | | | | |
| **Target** | **Clone** | **Vendor** | **Catalog number** | **Oligo** | **Fluorophore** | **Titer (dilution or ug/ml)** | **Exposure (ms)** |
| HLA-DR | E9R2Q | CST | 88471 | 65 | AlexaFluor-647 | 1 | 150ms |
| **Figure 2E: 8-plex RNAScope integrated with protein staining** | | | | | | | |
| **Target** | **Clone** | **Vendor** | **Catalog number** | **Oligo** | **Fluorophore** | **Titer (dilution or ug/ml)** | **Exposure (ms)** |
| NaKATP | D4Y7E | CST | 99833 | 25 | ATTO550 | 5 | 100ms |
| Ki67 | B56 | BD Bioscience | 556003 | 6 | AlexaFluor-647 | 1:32 | 100ms |

**Supplementary Table 1: Antibody Panels, conjugated oligonucleotides, and Imaging Conditions.** The table shows the antibodies used in **Fig. 1** and **Fig. 2**, the oligos they were conjugated to, the complementary fluorescent reporter oligo, and the exposure for imaging.

| Figure 1 | | | |
|---|---|---|---|
| **Target** | **HRP-oligo type** | **Manufacturer** | **Tyramine oligo Manufacturer** |
| Iba1 | HRP oligo | Biomers | GeneLink |
| CD45RA | HRP oligo | Biomers | GenScript |
| PD-L1 | HRP oligo | Biomers | GeneLink |
| CD20 | Biotin oligo | IDT | GeneLink |
| CD3 | HRP oligo | Biomers | GeneLink (oligo 1), GenScript (oligo 2) |
| Tox/Tox2 | HRP oligo | Biomers | GeneLink |
| CD11c | Biotin oligo | IDT | GeneLink (oligo 1 and 2) |
| Fas Ligand | HRP oligo | Biomers | GeneLink |
| **Figure 2A: EBV DNA Detection in PCNSL** | | | |
| **Target** | **HRP Type** | **Manufacturer** | **Tyramine oligo Manufacturer** |
| EBV DNA | RNAScope | ACDBio | Biomers |
| **Figure 2A: SARS-CoV-2 Spike RNA Detection in infected non-human primates** | | | |
| **Target** | **HRP Type** | **Manufacturer** | **Tyramine oligo Manufacturer** |
| SARS-CoV-2 Spike RNA | RNAScope | ACDBio | GenScript |
| **Figure 2B: Rescue of markers in RCC samples** | | | |
| **Target** | **HRP-oligo type** | **Manufacturer** | **Tyramine oligo Manufacturer** |
| CAIX | HRP oligo | Biomers | GenScript |
| CD4 | HRP oligo | Biomers | GeneLink |
| **Figure 2C: Conjugation-independent integration in colorectal carcinoma samples** | | | |
| **Target** | **HRP Type** | **Manufacturer** | **Tyramine oligo Manufacturer** |
| CD73 | Secondary Ab | Origene | GenScript |
| A2AR | Secondary Ab | Origene | GenScript |
| PD-L1 | Strep-HRP | ThermoFisher | GenScript |
| **Figure 2D: Integration of proximity ligation assays** | | | |
| **Target** | **HRP Type** | **Manufacturer** | **Tyramine oligo Manufacturer** |
| MHC II:CD4 PLA | Navinci PLA | Navinci | GeneLink |
| **Figure 2E: 8-plex RNAScope integrated with protein staining** | | | |
| **Target** | **HRP Type** | **Manufacturer** | **Tyramine oligo Manufacturer** |
| RPL28 | RNAScope | ACDBio | GeneLink |
| HPRT1 | RNAScope | ACDBio | GeneLink |
| B2M | RNAScope | ACDBio | GeneLink |
| POLR2A | RNAScope | ACDBio | GenScipt |
| TUBB | RNAScope | ACDBio | GeneLink |
| UBC | RNAScope | ACDBio | GeneLink |
| RPL5 | RNAScope | ACDBio | GenScipt |
| PPIB | RNAScope | ACDBio | GeneLink |

**Supplementary Table 2: HRP Sources and Tyramine Oligos.** The table shows different paths of HRP recruitment and the manufacturer of the corresponding HRP type. It also shows the tyramine oligonucleotide manufacturer.

| Figure 2A: EBV DNA probes | | | | | | |
|---|---|---|---|---|---|---|
| Target | Catalog number | LOT number | Fluorophore | Exposure | | |
| V-HHV4-06-C1 | 1324551-C1 | 24039A | ATTO550 | 100ms | | |
| **Figure 2A: SARS-CoV-2 Spike RNA probes** | | | | | | |
| **Target** | **Catalog number** | **LOT number** | **Fluorophore** | **Exposure** | | |
| nCoV2019-S-C1 | 848561-C2 | 24233A | AlexaFluor-647 | 50ms | | |
| **Figure 2E: 8-plex RNAScope Housekeeping gene probes** | | | | | | |
| | | | 8plex Run | | 4plex (Supp. Fig. 2) | |
| **Target** | **Catalog number** | **LOT number** | **Fluorophore** | **Exposure** | **Fluorophore** | **Exposure** |
| Hs-RPL28-C1 | 511521 | 24352B | AlexaFluor-647 | 200ms | DyLight 755 | 500ms |
| Hs-HPRT1-C2 | 310341-C2 | 24353C | AlexaFluor-647 | 250ms | AlexaFluor-647 | 5ms |
| Hs-B2M-C3 | 310161-C3 | 24353C | AlexaFluor-647 | 150ms | AlexaFluor-568 | 25ms |
| Hs-POLR2A-C4 | 310451-C4 | 24353C | AlexaFluor-647 | 150ms | AlexaFluor-488 | 5ms |
| Hs-TUBB-C1 | 588791 | 24352B | AlexaFluor-647 | 250ms | DyLight 755 | 500ms |
| Hs-UBC-C2 | 310041-C2 | 24353C | AlexaFluor-647 | 150ms | AlexaFluor-647 | 50ms |
| Hs-RPL5-C3 | 416361-C3 | 24353C | AlexaFluor-647 | 100ms | AlexaFluor-568 | 25ms |
| Hs-PPIB-C4 | 313901-C4 | 24353C | ATTO550 | 200ms | AlexaFluor-488 | 25ms |

**Supplementary Table 3: RNAScope Probes.** The table shows the RNAScope probes used in **Fig. 2** as well as associated imaging conditions. These probes are commercially available from ACDBio.
